## Supplementary Information - FIgure S1 for "Non-trivial dynamics in a model of glial membrane voltage driven by open potassium pores"

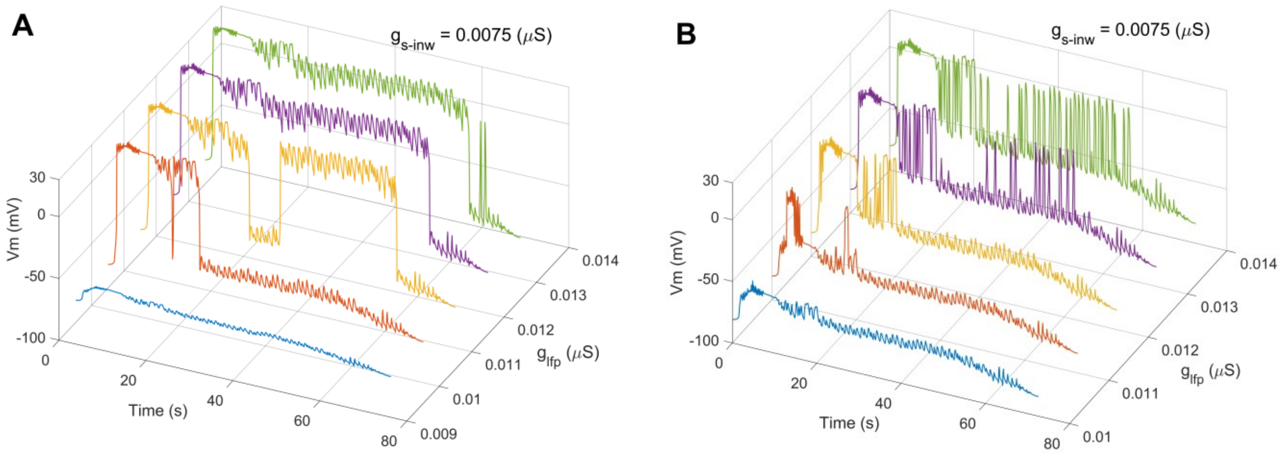


**Figure S1 – Glial switching on transient seizure-like perturbation, with Kir inward current decreased** – (**A**) Simulations of the full model $V_m$ response on electrographic seizure, Fig. 9, with fully abolished Kir outward $I_{res}$ current and increasing $g_{lfp}$ show plateau switching between downstate and upstate, corresponding to $V_r$ and $V_{dr}$. (**B**) The same simulation, with $I_{res}$ amplitude reduced to 15%, demonstrating a more irregular, erratic switching with a frequency following that of the clonic phase of the seizure signal. Presence of $I_{res}$ shifts both $V_r$ and $V_{dr}$ slightly more negative. A constant level of external current through gap junctions was kept at $I_{gjc} = 0.125\ nA$ and the slope of Kir inward conductance was decreased to $g_{s-inw} = 0.0075\ \mu S$ (for 18%). Compared to Fig. 10, decreased stabilizing effect of Kir inward current $I_{inw}$ results in more frequent bistability switching in both profiles.
